# Arabidopsis ubiquitin ligase PUB41 is a positive regulator of the ABA response

**DOI:** 10.1101/2024.04.07.588448

**Authors:** Avinash Sharma, Shalev Goldfarb, Dina Raveh, Dudy Bar-Zvi

**Affiliations:** Department of Life Sciences, Ben-Gurion University of the Negev, Beer-Sheva, Israel

**Keywords:** Arabidopsis, PUB E3 ubiquitin ligase, ABA signaling, dormancy, germination, drought

## Abstract

Seed germination is a highly controlled developmental process. Prevention of early germination is essential to avoid germination at a time that is not likely to allow the completion of the plant life cycle before entering unfavorable seasons. The plant hormone ABA is the key regulator of seed dormancy and inhibition of germination. ABA is also involved in the plant response to drought. Here we report on the involvement of AtPUB41, a U-BOX E3 ubiquitin ligase from *Arabidopsis thaliana*, in regulating ABA signaling, seed dormancy, germination, and drought resilience. *AtPUB41* is expressed in most plant vegetative and reproductive tissues. AtPUB41 protein is localized in the cytosol and nucleus. *pub41* T-DNA insertion mutants display reduced seed dormancy, and their germination is less inhibited by exogenous ABA than seeds of wild-type plants. *pub41* mutant plants are also hypersensitive to drought. ABA induces *AtPUB41* promoter activity and steady-state mRNA levels in the roots. Our data suggest that *AtPUB41* is a positive regulator of ABA signaling.

**Main Conclusion:** Arabidopsis ubiquitin ligase PUB41 is induced by ABA and acts as a positive regulator of ABA signaling by modulating seed dormancy, germination, and drought resilience.

## Introduction

As sessile organisms, plants are continuously exposed to fluctuating environments and have evolved mechanisms that modulate their growth and development in response to environmental cues. The plant hormone ABA is a key player in modulating these developmental processes, from the breaking of seed dormancy for germination reviewed in (Jones 2016), to the response to abiotic stresses, including drought (Ma et al. 2018).

ABA signaling pathways have been elucidated: ABA binds to the soluble receptors RCAR/PYR/PYL, which then bind and inhibit PP2C protein phosphatase. PP2C inactivation leads to activation of SnRK protein kinase that phosphorylates and activates downstream transcription factors leading to a global change in gene expression (Tang et al. 2015). Moreover, the ABA response mechanism involves a change in the cellular proteome, resulting from specific changes in protein translation and degradation. The Ubiquitin-Proteasome System (UPS) is a central mechanism for controlled protein degradation in eukaryotes including plants (Vierstra 2009) and has a major role in ABA signaling (Yu et al. 2016). For example, the protein levels of ABA receptors, PP2A protease, as well as various ABA-regulated transcription factors are controlled by ubiquitination-mediated degradation by the 26S proteasome (Yu et al. 2016).

A large part of the plant genome, about 5% in Arabidopsis, encodes UPS components reflecting the pivotal role of the UPS in maintaining proteome homeostasis. The majority of these genes (>1,400 in Arabidopsis) encodes E3s that specifically recognize and ubiquitinate target proteins (Vierstra 2009). In particular, the plant U-box (PUB) E3s have a major role in diverse biological processes, including development, immunity, and abiotic stress (reviewed by (Trujillo 2018, 2021; Vierstra 2009)). PUB E3s comprise a U-box domain that binds the E2 ubiquitin conjugating enzyme and several repeats of the Armadillo (ARM) motif that bind the protein targeted for proteasomal degradation (Adler et al. 2010).

Here, we present the role of the Arabidopsis U-box E3 ligase *PUB41* in ABA signaling by showing that ABA enhances the promoter activity of *AtPUB41* and the steady-state levels of this gene transcript, indicating that *AtPUB41* serves as a positive regulator of ABA signaling. We show that *AtPUB41* is important for seed dormancy, germination, and the drought response: seeds of *pub41* mutants showed reduced ability to maintain dormancy, and their germination was less sensitive to exogenous ABA. Furthermore, *pub41* T-DNA mutant plants are more resilient to drought than wild-type plants.

## Materials and methods

### Plant Material

Homozygous *Arabidopsis thaliana* ecotype Columbia was used in this study. T-DNA insertion lines *pub41-1* (SALK_099012) and *pub41-2* (SALK_142012) were obtained from the Arabidopsis Resource Center, Columbus, Ohio. The presence of T-DNA insertion was confirmed by Polymerase Chain Reaction (PCR) using T-DNA and gene-specific primers, further confirmed by Sanger sequencing of the PCR product. Primers are listed in Supplementary Table S1.

### Plant growth conditions

Plants were cultivated at 22°C, with a relative humidity of 50%, following a circadian cycle of 12 hours of light and 12 hours of darkness. Seeds were surface sterilized and imbibed at 4°C for at least 4 days before they were sown on solid 0.5 × Murashige and Skoog (MS) + 0.5% (w/v) sucrose or in pots as described previously (Adler et al. 2017). Post germination application of ABA was performed by incubating plate-grown 7-day-old seedlings for 8 h in the light at room temperature on Whatman No. 1 paper soaked in 0.5 x MS and with the indicated concentration of ABA.

### Seed germination

Surface-sterilized cold-treated seeds were sown on Petri plates containing 0.5 x MS and 0.7% agar, and the indicated ABA concentrations. Plates were incubated at 22 °C in a 12 h light/12 h dark regime. Radicle emergence was scored over 7 days.

### Drought tolerance

Plants were grown for 3 weeks in pots containing equal amounts of potting mix under non-stress conditions. Water was then withheld, and plant wilting and drying were followed daily. Upon wilting of the plants, the pots were rewatered, and plant survival was visually scored 4 days later.

### Transcript levels

RNA isolation, cDNA synthesis, primer design, and RT-qPCR assays for determining relative steady-state transcript levels were performed as previously described (Maymon et al. 2022). Primers are listed in Supplementary Table S1.

### Plasmid construction and plant transformation

The 35S::PUB41-eGFP plasmid was constructed by amplifying the DNA sequence encoding the full-length PUB41 with primers containing linkers with the *Nco*I and *Pst*I restriction sites. This sequence was subcloned into the respective restriction sites of the pGA-eGFP3 vector (Maymon et al. 2022) in frame with the sequence encoding eGFP.

To construct *AtPUB41::GUS* expressing lines, a 1440 bp DNA fragment upstream to the start codon of the PUB41 gene was amplified using the primers listed in Supplemental Table S1 and genomic DNA of wild-type Arabidopsis plants. The amplified *AtPUB41* promoter DNA fragment was sub-cloned into the *Pst*I and *Eco*RI restriction sites of pCAMBIA 1391Z upstream to the GUS encoding sequence.

Plasmids were verified by DNA sequencing and introduced into *Agrobacterium tumefaciens* GV3101, and wild-type Arabidopsis plants were transformed by the floral dip method (Clough and Bent 1998). Homozygous transformants were isolated by selection on plates containing growth medium supplemented with hygromycin. Primers are listed in Supplementary Table S1.

### GUS staining

Plant tissues, upon treatment, were fixed into acetone chilled at -20 °C, and GUS staining was performed as described (Jefferson 1987). Images were taken using a dissecting microscope equipped with a Dino-Eye: AM7025X Edge Series 5MP Eyepiece Camera.

### Statistical analysis

All experiments were carried out in at least 3 biological repeats. Data are expressed as average ± SE. Statistical significance was determined by using Tukey’s HSD post hoc test and the Student’s t-test.

## Results and Discussion

### Domain structure and cellular localization of PUB41

The Arabidopsis *PUB41* gene (At5G62560) is an intronless gene that encodes a 559 amino-acid long E3 of the Plant-U-box (PUB) family. AtPUB41 has ubiquitin ligase activity (Wiborg et al. 2008). Motif analysis of the amino acid sequence of AtPUB41 indicates a U-box motif at residues 30 to 104, and a cluster of 5 ARM motifs located between residues 266-305, 307-346, 348-388, 390-427 and 428-472 (Fig. 1A).

**Fig. 1.**
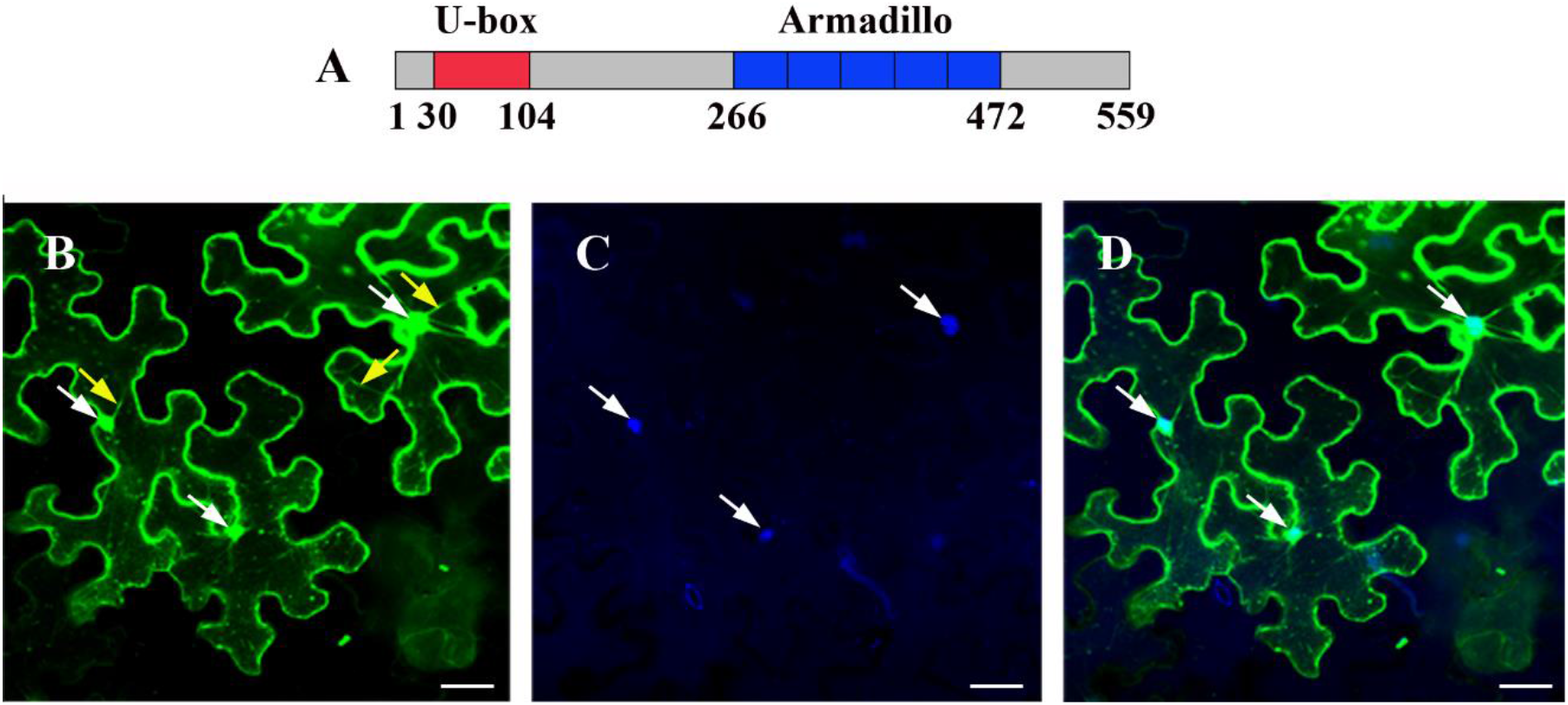
Domain organization and subcellular localization of AtPUB41. A. Diagram presenting the linear amino-acid sequence of AtPUB41 and functional motifs. Numbers represent the respective amino acids. B-D. Subcellular localization of AtPUB41. Leaves of *N. benthamiana* were infiltrated with Agrobacteria harboring *35S::AtPUB41-eGFP*. Lower epidermis peels were prepared and counterstained with the DNA marker DAPI, and fluorescence GFP (B) and DAPI (C) were analyzed using confocal microscopy. (D), merged image of (B) and (C). White and yellow arrows point at nuclei and transvacuolar cytoplasmic strands, respectively. Scale bar = 50 μm.

To assess the subcellular localization of the AtPUB41 protein, we transiently expressed AtPUB41-eGFP in leaves of *Nicotiana benthamiana*. Lower epidermis peels were counterstained with the DNA binding stain DAPI (4′,6-diamidino-2-phenylindole), and cells were examined by confocal fluorescence microscopy. AtPUB41-eGFP was detected in the nucleus and the cytoplasm (Fig. 1). The cytoplasmic localization of AtPUB41 is supported by its presence in transvacuolar cytoplasmic strands (Fig. 1B), and the nuclear localization is demonstrated by colocalization of the AtPUB41-GFP signal with that of the DNA fluorescent stain (Fig. 1D). The NLS mapper (Kosugi et al. 2009) supports the nuclear import of AtPUB41, indicating three presumptive nuclear localization signals (NLSs) at residues 3-32, 165-194, and 256-287, with scores of 3.7, 3.5, and 3.2, respectively, supporting our experimental demonstration of its nuclear localization.

### Promoter activity of *AtPUB41*

Only limited expression data for At*PUB41* is available as the AT5G62560 locus was not included in the Affymetrix ATH1 microarrays. Therefore to examine the activity of the *AtPUB41* promoter, we transformed Arabidopsis plants with a plasmid encoding the β-glucuronidase (GUS) reporter gene driven by the *AtPUB41* promoter (*AtPUB41::GUS*), comprising 1440 bp upstream of the translation start codon. Plants of three independent homozygous lines were analyzed. Under controlled optimal growth conditions, the *AtPUB41* promoter was predominantly active in cotyledons, leaf vascular tissue, petioles, hydathodes, axillary buds, stems, and roots (Fig. 2A). High promoter activity was observed in embryos (Fig. 2B) and endosperm (Fig. 2C) of imbibed seeds. *AtPUB41* promoter activity was detected in fully expanded leaves (Fig. 2E-G), where, in addition to vascular tissue, notable expression was detected in trichomes and stomata guard cells (Fig. 2E-G). The *AtPUB41* promoter activity was also present in the root hairs of the collet (Fig. 2D) and root maturation zone (Fig. 2I). In mature plants, promoter activity was detected in sepals, pistils, anther filaments, and mature siliques (Fig. 2J-K). In contrast, promoter activity was not detected in the root elongation zone, mature anthers, and petals (Fig. 1D, H & J). Moreover, our RT-qPCR analysis of 10-day-old seedlings revealed a higher expression of PUB41 in roots than in shoots, indicating a role for PUB41 in root tissue (Fig. 2L).

**Fig. 2.**
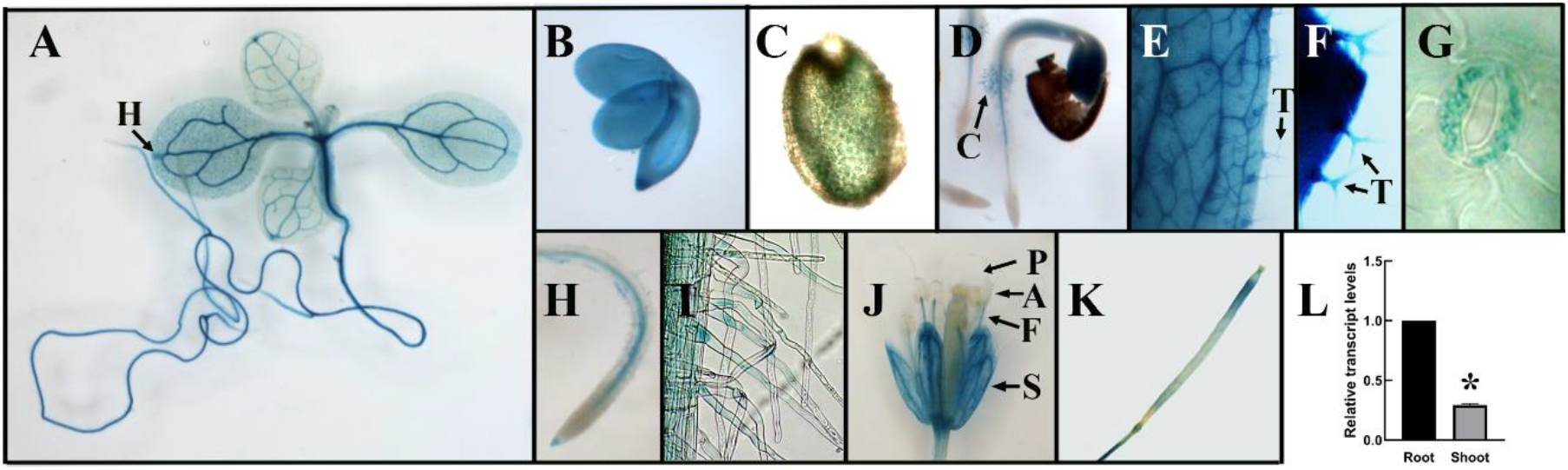
Tissue specificity of *AtPUB41* expression. (A-K) Arabidopsis plants expressing GUS reporter gene from the *AtPUB41* promoter stained for GUS activity. A. 8-day old seedling. Arrow marks hydathode (H). B. Embryo rescued from imbibed seed. C. Endosperm cells in imbibed seed. D. 36-hour-old Arabidopsis seedling. Arrow points to promoter activity in collet region root hairs (C). E & F. Sections of fully expanded leaves. Arrows point tot trichomes (T). G. Stomata. H. Root tip and root meristem zone of the primary root. I. Mature root hairs of the maturation zone. J. Open flower: A, anther; F, anther-filament; P, petal; S, sepal. K. Silique. L. Relative PUB41 transcript levels. The roots and shoots of ten-day-old wild-type seedlings grown on 0.5 x MS without supplements were harvested, RNA was isolated, and PUB41-transcript levels were determined, by RT-qPCR. Data showing mean ± SE of three repeats. Asterisks denote statistical significance as determined by a two-tailed paired student’s t-test (*p ≤ 0.001).

### *pub41* T-DNA insertion lines

Two *pub41* T-DNA insertion mutants were obtained from the Arabidopsis stock center, Columbus, Ohio. SALK_099012C and SALK_142330 are designated as *pub41-1* and *pub41-2*, respectively (Fig. 3A). The mutants were verified by PCR analysis using gene-specific and T-DNA border primers (Supplemental Table 1), as described earlier (O’Malley et al. 2015) Only the *pub41-1* and *pub41-2* lines yielded a product of ∼ 500 bp with the LBb1.3 and RP primers (Fig. 3B). The amplified PCR product was subsequently used for Sanger sequencing to identify the precise location of the T-DNA insertion. The sequencing results revealed that the T-DNA inserted within the protein-encoding region (+1505 bp downstream from the start codon) in *pub41-1*, and within the 5’ untranslated region (UTR) (−69 bp upstream to the start codon in the *pub41-2* mutant (Fig. 3A). RT-qPCR analysis further confirmed that transcript levels of *AtPUB41* in both mutants are less than 18% that of the wild-type plants (Fig. 3C).

**Fig. 3.**
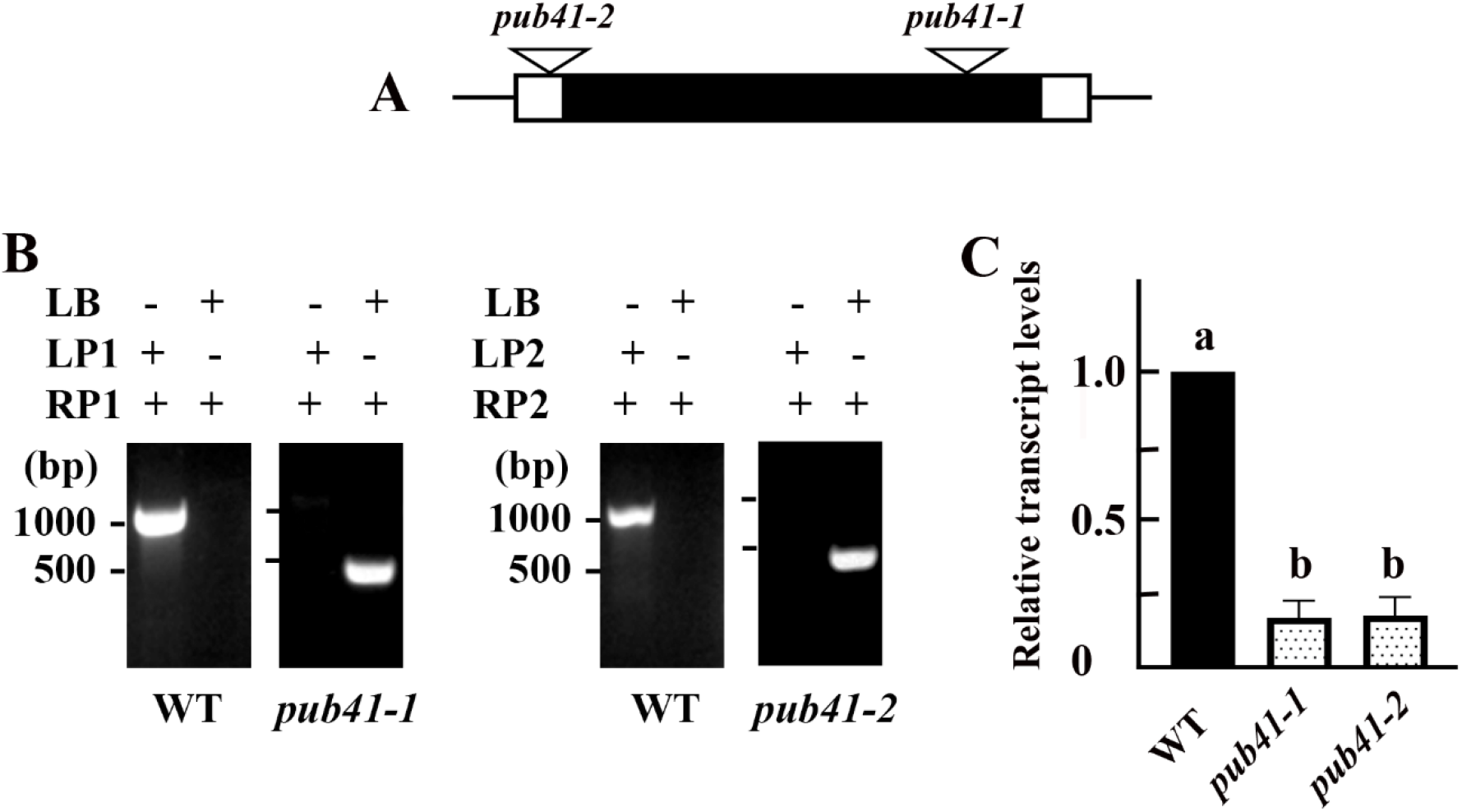
Molecular characterization of *AtPUB41* T-DNA insertion lines. (A) Diagram of the *AtPUB41* gene. Transcribed sequences are shown as boxes, where the black region marks the protein-encoding sequence, and the white regions mark 5’ and 3’ UTRs. T-DNA insertion sites of the respective mutants are marked by triangles. (B) Verification of the T-DNA insertion loci by PCR. Genomic DNA from the indicated genotypes was amplified using the T-DNA left border (LB) primer and site-specific left and right primers (LP1, LP2, and RP1, RP2) as indicated. (C) Steady-state levels of *AtPUB41* transcripts in 5-day old seedlings of wild-type (WT) and *pub41* mutants. The data represent the mean ± SE of three independent experiments. Bars with different letters represent statistically different values by ordinary one-way ANOVA, Tukey’s multiple comparison test (p< 0.001).

### *pub41* mutants are hypersensitive to drought

Our original screen to identify UPS-associated genes that affect drought resilience led to the identification of 3 PUB genes, *AtPUB8, AtPUB46*, and *AtPUB41*. The involvement of each of these genes was confirmed using a single T-DNA insertion mutant (Adler 2011). To further confirm the drought-hypersensitive phenotype of the *pub41* mutant and to rule out the possibility that the drought hypersensitivity resulted from a positional effect, we repeated the water-withholding experiment using two independent *pub41* mutants. Fig. 4 shows that both *pub41-1* and *pub41-2* mutants are hypersensitive to drought, and under the protocol applied, they failed to recover from water-withholding stress, in contrast to the wild-type plants. These results show that *AtPUB41* is essential for Arabidopsis drought stress resilience.

**Fig. 4.**
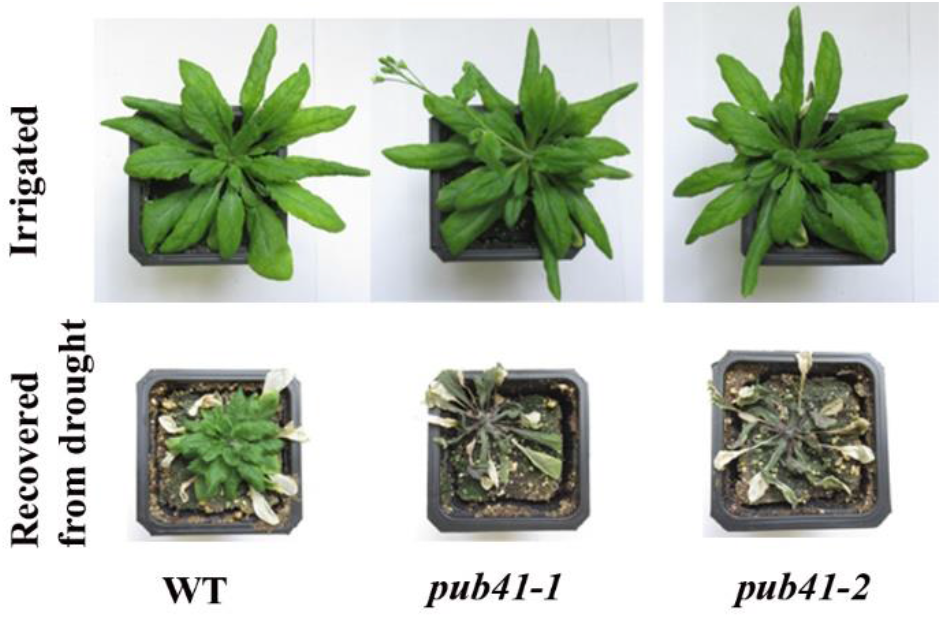
*pub41* mutant plants are more sensitive to drought than wild-type plants. Wild-type (WT) and *pub41* mutant plants were grown in pots for six weeks before water was withheld for 12 days. Water was supplied only after the plants were fully wilted and the plants were imaged 4 days later in the recovery period.

Ubiquitin ligases belonging to the PUB family are recognized for their involvement in the plant response to abiotic and biotic stresses (Trujillo 2018, 2021; Vierstra 2009). Interestingly, pub mutants reported to be affected in drought resilience were both hyper- or hypo-sensitive: single gene mutants of the Arabidopsis paralogs *AtPUB46* and *AtPUP48* (Adler et al. 2017), and the *Atpub8* mutant (Adler 2011) are drought hypersensitive whereas Arabidopsis *Atpub18, Atpub19, Atpub23*, and *Atpub24* mutants exhibit increased drought tolerance compared with wild-type plants (Seo et al. 2012). Also, the rice OsPUB7 knockout mutant and the *Ospub41* suppression mutant, and *Ubi:RNAi-OsPUB41* knock-down lines showed enhanced survival to drought (Kim et al. 2023; Seo et al. 2021). (Despite sharing a common gene name *OsPUB41* is not orthologous to *AtPUB41*, but shows the highest similarity to *AtPUB20* and *AtPUB21*).

### Seed germination of *pub41* mutants shows reduced ABA inhibitory effects

In addition to drought, ABA is also the primary hormone that ensures seed dormancy and represses germination (Bewley et al. 2013; Ma et al. 2018). We therefore examined the impact of ABA on germination of seeds of the *pub41* mutants. Under non-stress conditions, the germination of *pub41* mutants was similar to that of wild-type plants (Fig. 5A). However, the germination of seeds of the *pub41-1* and *pub41-2* mutant lines displayed reduced sensitivity to ABA inhibition than those of the wild-type plants (Fig. 5B). For example, 52 h following plating on growth medium containing 2.5 μM ABA, 25.3% and 22.7% of seeds of the *pub41-1* and *pub41-2* mutants germinated compared with only 6.7% of the wild-type seeds.

**Fig. 5.**
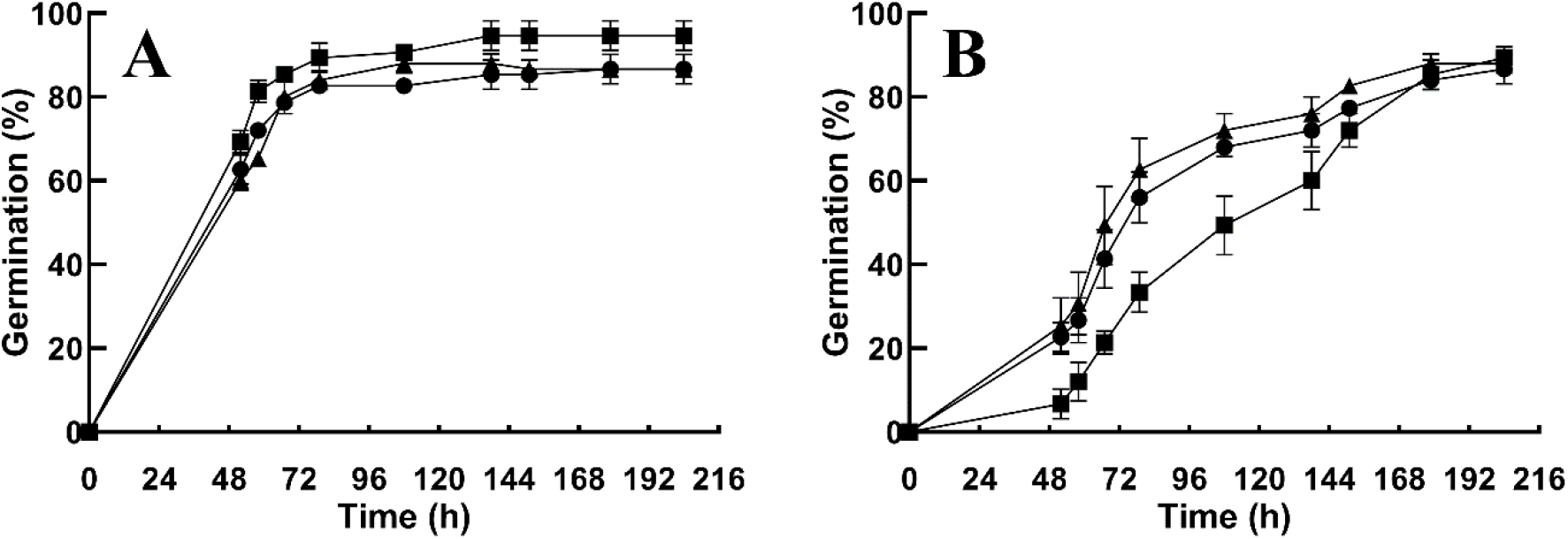
Germination of *pub41* mutants is less sensitive to ABA inhibition. Surface sterilized cold-treated seeds of the wild-type (WT) (▪), *pub41-1* (▲), and *pub41-2* (•) genotypes were plated on agar-solidified 0.5 X MS 0.5% sucrose medium without (A) or with 2.5 μM ABA (B). Germination was scored at the indicated times. Data represent means ± SE of three independent biological repeats.

### Seeds of *pub41* mutants show reduced dormancy

The rate of seed germination depends upon the speed of ABA degradation upon imbibition (Preston et al. 2009). Freshly collected Arabidopsis seeds display suppressed germination rates. Seed dormancy can be relieved with storage time of dry seeds, and can be reversed by stratification of the imbibed seeds at any time point (Bewley et al. 2013). ABA is the primary hormone in maintenance of dormancy, and gibberellins (GA) are the key players in germination (Gianinetti 2023). Thus, the ABA to GA ratio determines the fate of germination. Here we assayed germination using freshly collected non-stratified seeds. *pub41-1* and *pub41-2* seeds germinated faster than the corresponding wild-type seeds, indicating that in the absence of functional PUB41 there is a reduction in the ability to maintain dormancy (Fig. 6A). As expected, cold treatment of all tested genotypes broke dormancy and resulted in faster and similar kinetics of germination (Fig. 6B), supporting our interpretation that the differences in germination of the non-treated seeds of the *pub41* mutants and wild-type plants result from reduced dormancy of the *pub41* mutants. Interestingly, a significant quantity of ABA is produced by a single layer of endosperm surrounding the embryo during germination (Karssen et al. 1983). This is compatible with the high PUB41 promoter activity in embryos and endosperm we observed above (Fig. 2B & C).

**Fig. 6.**
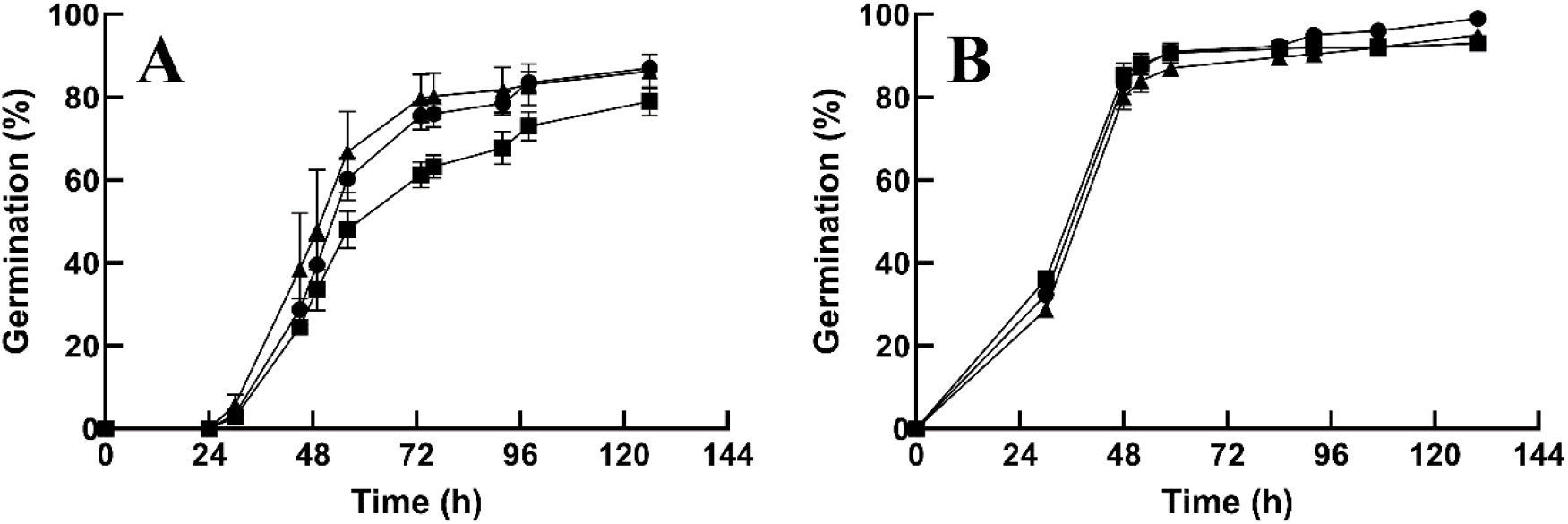
Germination of newly collected seeds. Freshly harvested, surface sterilized seeds of the wild-type (WT) (▪), *pub41-1* (▲), and *pub41-2* (•) genotypes were plated on agar solidified 0.5 X MS 0.5% sucrose medium without pre-treatment (A) or after stratification (B). Germination was scored at the indicated times. Data represent means ± SE of three independent biological repeats.

### ABA induces the expression of *AtPUB41*

Although ABA signaling results in a global change of gene expression (Rock 2000), the hormone does not affect the expression of genes involved in the response to ABA itself (for example, see (Pardo-Hernández et al. 2024)). We thus treated seedlings with ABA and assayed the effect on *AtPUB41* expression. ABA treatment increases the steady-state levels of *AtPUB41* transcript in the root by 250% (Fig. 7A). In histochemical staining of plants expressing *AtPUB41::GUS* (Fig. 7B - E), we observed that ABA treatment enhanced *AtPUB41* promoter activity in the roots. Close to the root tip, *AtPUB41* promoter activity was present in all cell types whereas in the upper root zone it was primarily in the vascular system (Fig. 7D & E). Thus, *AtPUB41* behaves like a typical ABA-responsive gene whose expression is increased by this plant hormone.

**Fig. 7.**
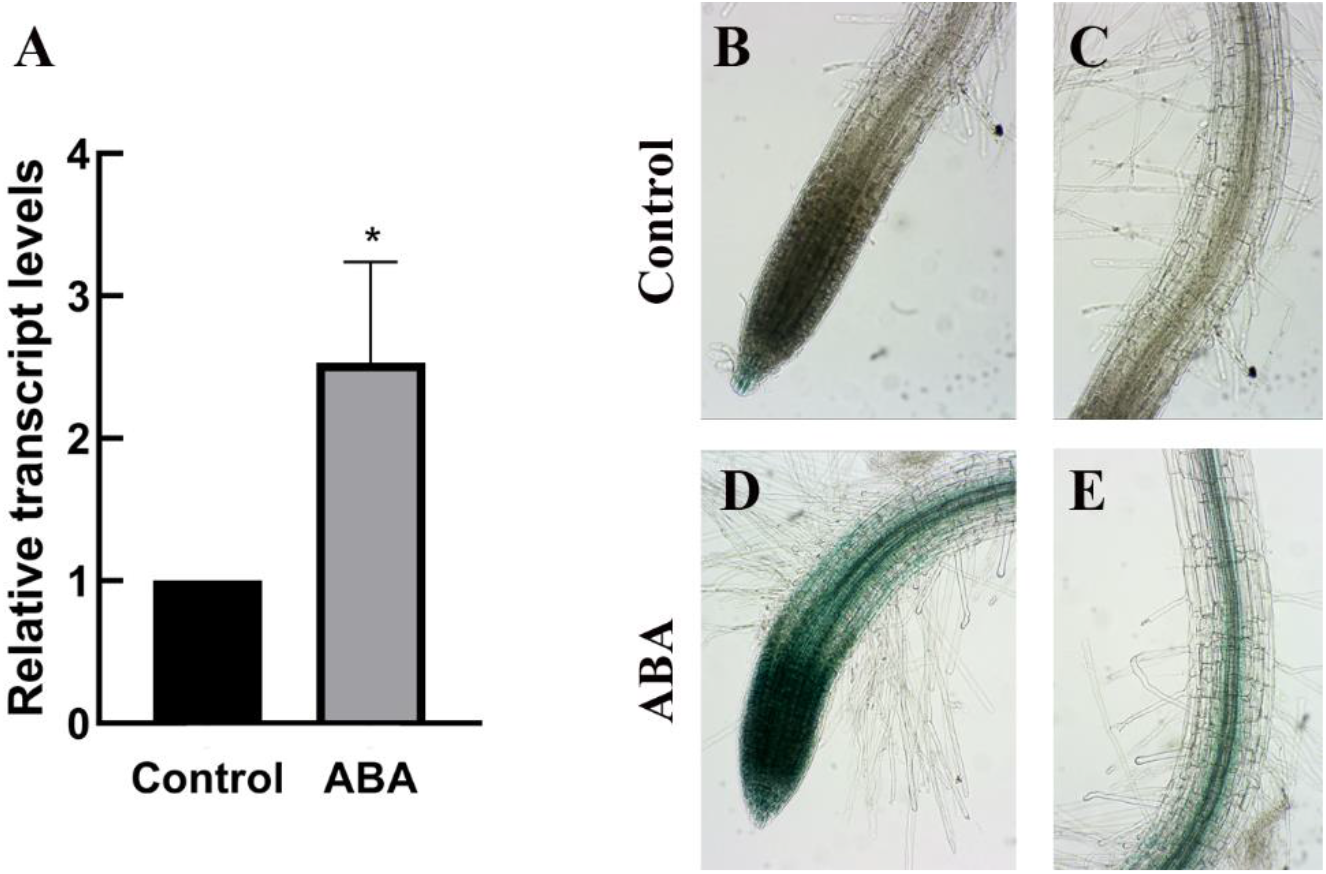
ABA induces *AtPUB41* expression. (A) RT-qPCR results showing *AtPUB41* transcript levels in roots of ten-day-old wild-type plants treated for 8 h without or with 10 µM ABA. Data show mean ± SE of three biological repeats. Asterisks denote statistical significance as determined by a two-tailed paired student’s t-test (P ≤ 0.05). (B-E) One week old *AtPUB41::GUS* Arabidopsis seedlings were treated for 8 h without (B-C) or with 10 µM ABA (D-E), followed by histological staining for GUS activity. (B & D) primary root tips; (D & E) differentiation zone of the primary roots.

### *AtPUB41* is a positive regulator of ABA signaling

Our findings show that *AtPUB41* is a positive regulator of ABA signaling. Germination of *pub41* mutants was less inhibited by ABA (Fig. 5) and the seeds displayed reduced dormancy (Fig. 6). Dormancy was shown to be regulated by ABA signaling rather than ABA levels in the seeds (Footitt et al. 2011). Most PUB genes involved in the plant response to environmental stress function as negative regulators, i.e., the mutants are more resilient to stress than the WT, and overexpressing the respective genes often results in hypersensitivity. These include: *AtPUB18, AtPUB19, AtPUB22* and *AtPUB23* that function as negative regulators of ABA-mediated drought responses (Cho et al. 2008; Seo et al. 2012). AtPUB12/13 degrade the ABA co-receptor ABI1 (Kong et al. 2015), and *Atpub12 Atpub13* mutants are less sensitive to ABA. Steady-state mRNA levels of the Ubox E3 AtCHIP increased following exposure to heat, cold, and salt treatments, and overexpression of this gene increased plant sensitivity to temperature stress (Yan et al. 2003).

The soybean *GmPUB8* (Wang et al. 2016) and wheat *TaPUB26* (Wu et al. 2020) are negative regulators of the drought stress response *AtPUB11* negatively regulates drought through degrading a receptor-like kinase (Chen et al. 2021). *AtPUB22/23* negatively regulates drought tolerance by targeting the ABA receptor (Zhao et al. 2017). Rice *OsPUB41*, which is a non-ortholog of *AtPUB41*, is a negative regulator of drought (Seo et al. 2021). *AtPUB10* negatively regulates the ABA response (Seo et al. 2019. AtPUB25 and 26 negatively regulate immune response and disease resistance {Wang, 2018 #143). *AtPUB30* negatively regulates salt tolerance (Zhang et al. 2017).

In contrast, fewer PUBs act as positive regulators of the abiotic stress response: We previously showed that the paralogous genes *AtPUB46* and *AtPUB48* are positive regulators of the drought response, with single mutants being hypersensitive to drought (Adler et al. 2017). Furthermore, *AtPUB25* and *AtPUB26* positively regulate freezing tolerance (Wang et al. 2019), and *OsPUB67* is a positive regulator of drought tolerance (Qin et al. 2020).

PUB E3s have been implicated for their role in seed germination. *Atpub9*-KO mutants display hypersensitivity to ABA during seed germination (Samuel et al. 2008); suppression of the Arabidopsis *AtPUB30* results in decreased inhibition of germination by NaCl (Hwang et al. 2015); *AtPUB18* and *AtPUB19* genes are involved in salt inhibition of germination, with germination of double mutant plants being less sensitive to NaCl (Bergler and Hoth 2011).

*pub41* mutants are drought hypersensitive at the rosette stage and are less sensitive to ABA inhibition of germination than wild-type plants (Figs. 4 & 5). Similar phenotypes have been reported for other ubiquitin ligases, for example, single gene mutants of the paralogs Arabidopsis C3H2C3-type RING E3 ubiquitin ligases *Arabidopsis ABA-insensitive RING protein* (*AIRP*) *AtAIRP1, AtAIRP2, AtAIRP3* (Ryu et al. 2010; Cho et al. 2011; Kim and Kim 2013) and *SALT- AND DROUGHT-INDUCED RING FINGER1* (*SDIR1*) (Zhang et al. 2007). In addition, mutants of the Arabidopsis F-box E3 ligases *ABA-RESPONSE KELCH PROTEIN 1* (*ARKP1*) (Li et al. 2016), *F-BOX OF FLOWERING 2* (*FOF2*) (Qu et al. 2020), *EID1-like protein 3* (*EDL3*) (Koops et al. 2011) also exhibit this phenotype. Like PUB41, all these genes, except SDIR1, are induced by ABA. AtPUB41 is localized in both the cytosol and nucleus (Fig. 1). SDIR1 is localized in both cytosol and nucleus (Zhang et al. 2007), whereas AIRP1-3 are cytosolic (Ryu et al. 2010; Cho et al. 2011; Kim and Kim 2013), and EDL3 and FOF2 are nuclear (Koops et al. 2011; Qu et al. 2020), suggesting that the phenotype shared by all mutants may result from the activity of the respective E3 in either or both these subcellular compartments.

*AtPUB41* belongs to a 4 member-subfamily comprising *AtPUB38-AtPUB41* (Wiborg et al. 2008). *AtPUB40* was shown to mediate the root degradation of BZR1, a brassinosteroid-responsive transcription factor (Kim et al. 2019). Roots of the triple *pub39 pub40 pub41* mutant accumulated higher levels of BZR1 than WT. Unfortunately, Kim et al. (Kim et al. 2019) did not study the phenotype of the *Atpub41* single gene mutant, or *AtPUB41* overexpressor; thus, the role of *AtPUB41* in the brassinosteroid root signaling pathway remains unclear. Our data clearly show that *AtPUB41* is a positive regulator of ABA signaling, affecting seed germination and the drought response. The AtPUB41 target(s) within the ABA signaling pathway remain to be identified.

## Author Contribution Statement

DBZ, SG and AS conceived and designed research. AS and SG conducted experiments. DBZ,DR and AS wrote the manuscript. All authors read and approved the manuscript.

## Data Availability

All data supporting the findings of this study are available within the paper and its Supplementary Information.

## Acknowledgments

DBZ is the incumbent of The Israel and Bernard Nichunsky Chair in Desert Agriculture, Ben-Gurion University of the Negev.

---

**Supplementary Table S1.**
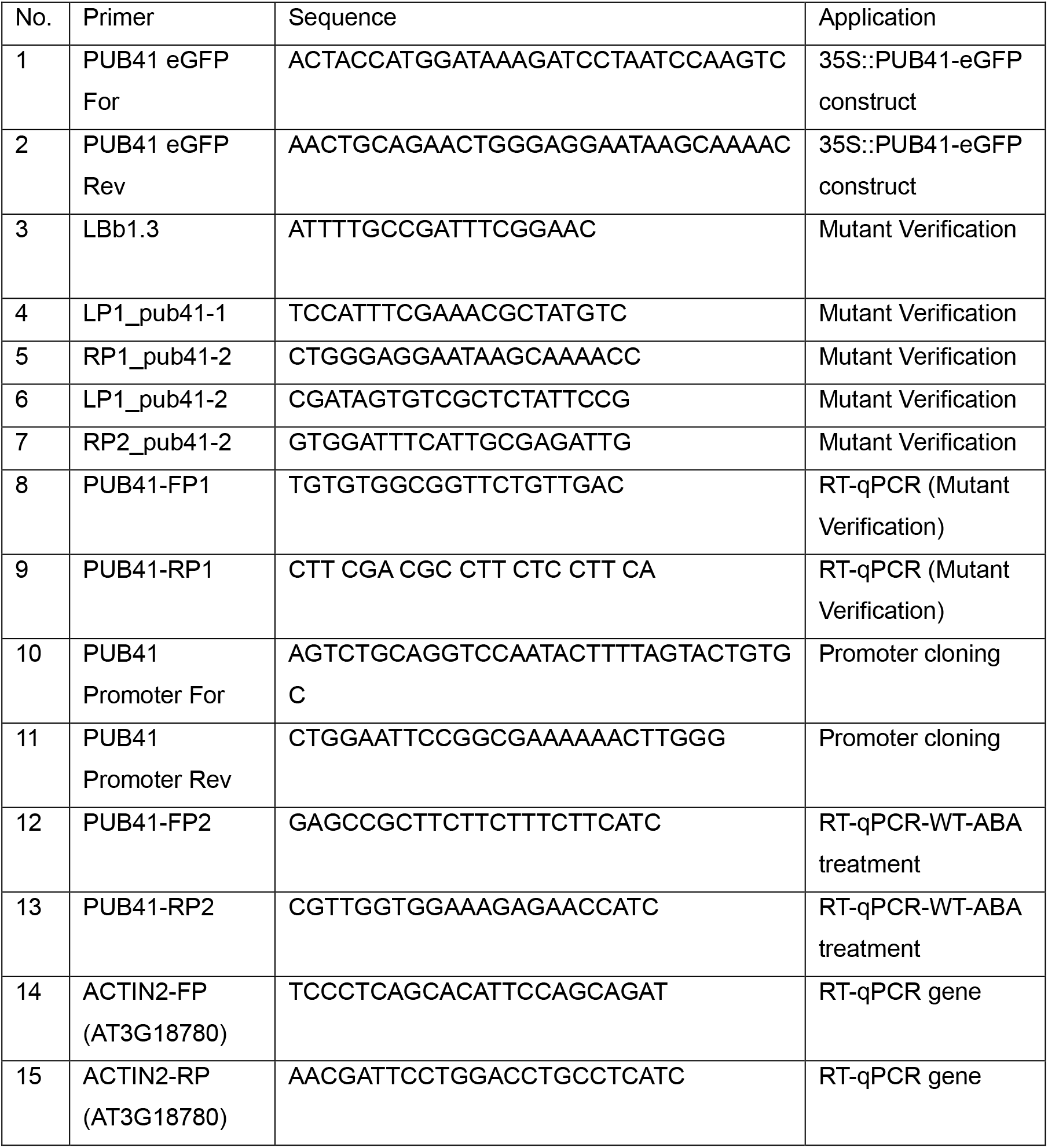
Primers used in this study.

## References

Adler G (2011) Plant response to environmental stresses: Expression regulation of BvNHX1 and the role of AtPUB proteins. Dissertation, Ben-Gurion University of the Negev, Beer-Sheva, Israel

Adler G, Konrad Z, Zamir L, Mishra AK, Raveh D, Bar-Zvi D (2017) The Arabidopsis paralogs, PUB46 and PUB48, encoding U-box E3 ubiquitin ligases, are essential for plant response to drought stress. BMC Plant Biol 17:8. 10.1186/s12870-016-0963-5

Bergler J, Hoth S (2011) Plant U-box armadillo repeat proteins AtPUB18 and AtPUB19 are involved in salt inhibition of germination in Arabidopsis. Plant Biol 13:725–730. 10.1111/j.1438-8677.2010.00431.x

Bewley JD, Bradford KJ, Hilhorst HWM, Nonogaki H (2013) Dormancy and the control of germination. In: Bewley JD, Bradford KJ, Hilhorst HWM, Nonogaki H (eds) Seeds: Physiology of Development, Germination and Dormancy, 3rd Edition. Springer New York, New York, NY, pp 247–297. 10.1007/978-1-4614-4693-4_6

Chen X, Wang T, Rehman AU, Wang Y, Qi J, Li Z, Song C, Wang B, Yang S, Gong Z (2021) Arabidopsis U-box E3 ubiquitin ligase PUB11 negatively regulates drought tolerance by degrading the receptor-like protein kinases LRR1 and KIN7. J Integr Plant Biol 63:494–509. 10.1111/jipb.13058

Cho SK, Ryu MY, Seo DH, Kang BG, Kim WT (2011) The Arabidopsis RING E3 ubiquitin ligase AtAIRP2 plays combinatory roles with AtAIRP1 in abscisic acid-mediated drought stress responses. Plant Physiol 157:2240–2257. 10.1104/pp.111.185595

Cho SK, Ryu MY, Song C, Kwak JM, Kim WT (2008) Arabidopsis PUB22 and PUB23 are homologous U-box E3 ubiquitin ligases that play combinatory roles in response to drought stress. Plant Cell 20:1899–1914. 10.1105/tpc.108.060699

Clough SJ, Bent AF (1998) Floral dip: a simplified method for Agrobacterium-mediated transformation of Arabidopsis thaliana. Plant J 16:735–743. 10.1046/j.1365-313x.1998.00343.x

Footitt S, Douterelo-Soler I, Clay H, Finch-Savage WE (2011) Dormancy cycling in Arabidopsis seeds is controlled by seasonally distinct hormone-signaling pathways. Proc Natl Acad Sci USA 108:20236–20241. 10.1073/pnas.1116325108

Gianinetti A (2023) A travel through landscapes of seed dormancy. Plants12:3963. 10.3390/plants12233963

Hwang JH, Seo DH, Kang BG, Kwak JM, Kim WT (2015) Suppression of Arabidopsis AtPUB30 resulted in increased tolerance to salt stress during germination. Plant Cell Rep 34:277–289. 10.1007/s00299-014-1706-4

Jefferson RA (1987) Assaying chimeric genes in plants: The GUS gene fusion system. Plant Mol Biol Rep 5:387–405. 10.1007/BF02667740

Jones AM (2016) A new look at stress: abscisic acid patterns and dynamics at high-resolution. New Phytol 210:38–44. 10.1111/nph.13552

Karssen CM, Brinkhorst-van der Swan DL, Breekland AE, Koornneef M (1983) Induction of dormancy during seed development by endogenous abscisic acid: studies on abscisic acid deficient genotypes of Arabidopsis thaliana (L.) Heynh. Planta 157:158–165. 10.1007/bf00393650

Kim EJ, Lee SH, Park CH, Kim SH, Hsu CC, Xu S, Wang ZY, Kim SK, Kim TW (2019) Plant U-Box40 mediates degradation of the brassinosteroid Responsive transcription factor BZR1 in Arabidopsis roots. Plant Cell 31:791–808. 10.1105/tpc.18.00941

Kim JH, Kim WT (2013) The Arabidopsis RING E3 ubiquitin ligase AtAIRP3/LOG2 participates in positive regulation of high-salt and drought stress responses. Plant Physiol 162:1733–1749. 10.1104/pp.113.220103

Kim MS, Ko SR, Jung YJ, Kang KK, Lee YJ, Cho YG (2023) Knockout mutants of OsPUB7 generated using CRISPR/Cas9 revealed abiotic stress tolerance in rice. Int J Mol Sci 24:5338. 10.3390/ijms24065338

Kong LY, Cheng JK, Zhu YJ, Ding YL, Meng JJ, Chen ZZ, Xie Q, Guo Y, Li JG, Yang SH, Gong ZZ (2015) Degradation of the ABA co-receptor ABI1 by PUB12/13 U-box E3 ligases. Nat Commun 6:8630. 10.1038/ncomms9630

Koops P, Pelser S, Ignatz M, Klose C, Marrocco-Selden K, Kretsch T (2011) EDL3 is an F-box protein involved in the regulation of abscisic acid signalling in Arabidopsis thaliana. J Exp Bot 62:5547–5560. 10.1093/jxb/err236

Kosugi S, Hasebe M, Tomita M, Yanagawa H (2009) Systematic identification of cell cycledependent yeast nucleocytoplasmic shuttling proteins by prediction of composite motifs. Proc Natl Acad Sci USA 106:10171–10176. 10.1073/pnas.0900604106

Li Y, Liu ZB, Wang JM, Li XF, Yang Y (2016) The Arabidopsis kelch repeat F-box E3 ligase ARKP1 plays a positive role for the regulation of abscisic acid signaling. Plant Mol Biol Rep 34:582–591. 10.1007/s11105-015-0942-2

Ma YL, Cao J, He JH, Chen QQ, Li XF, Yang Y (2018) Molecular mechanism for the regulation of ABA homeostasis during plant development and stress responses. Int J Mol Sci 19:3643. 10.3390/ijms19113643

Maymon T, Eisner N, Bar-Zvi D (2022) The ABCISIC ACID INSENSITIVE (ABI) 4 transcription factor Is stabilized by stress, ABA and phosphorylation. Plants 11:2179. 10.3390/plants11162179

O’Malley RC, Barragan CC, Ecker JR (2015) A user’s guide to the Arabidopsis T-DNA insertion mutant collections. Methods Mol Biol 1284:323–342. 10.1007/978-1-4939-2444-8_16

Pardo-Hernández M, Arbona V, Simón I, Rivero RM (2024) Specific ABA-independent tomato transcriptome reprogramming under abiotic stress combination. Plant J 117:1746–1763. 10.1111/tpj.16642

Preston J, Tatematsu K, Kanno Y, Hobo T, Kimura M, Jikumaru Y, Yano R, Kamiya Y, Nambara E (2009) Temporal expression patterns of hormone metabolism genes during imbibition of Arabidopsis thaliana seeds: a comparative study on dormant and non-dormant accessions. Plant Cell Physiol 50:1786–1800. 10.1093/pcp/pcp121

Qin Q, Wang Y, Huang L, Du F, Zhao X, Li Z, Wang W, Fu B (2020) A U-box E3 ubiquitin ligase OsPUB67 is positively involved in drought tolerance in rice. Plant Mol Biol 102:89–107. 10.1007/s11103-019-00933-8

Qu L, Sun M, Li X, He R, Zhong M, Luo D, Liu X, Zhao X (2020) The Arabidopsis F-box protein FOF2 regulates ABA-mediated seed germination and drought tolerance. Plant Sci 301:110643. 10.1016/j.plantsci.2020.110643

Rock CD (2000) Tansley Review No. 120. New Phytol 148:357–396. 10.1046/j.1469-8137.2000.00769.x

Ryu MY, Cho SK, Kim WT (2010) The Arabidopsis C3H2C3-type RING E3 ubiquitin ligase AtAIRP1 is a positive regulator of an abscisic acid-dependent response to drought stress. Plant Physiol 154:1983–1997. 10.1104/pp.110.164749

Samuel MA, Mudgil Y, Salt JN, Delmas F, Ramachandran S, Chilelli A, Goring DR (2008) Interactions between the S-domain receptor kinases and AtPUB-ARM E3 ubiquitin ligases suggest a conserved signaling pathway in Arabidopsis. Plant Physiol 147:2084–2095. 10.1104/pp.108.123380

Seo DH, Lee A, Yu SG, Cui LH, Min HJ, Lee SE, Cho NH, Kim S, Bae H, Kim WT (2021) OsPUB41, a U-box E3 ubiquitin ligase, acts as a negative regulator of drought stress response in rice (Oryza Sativa L.). Plant Mol Biol 106:463–477. 10.1007/s11103-021-01158-4

Seo DH, Ryu MY, Jammes F, Hwang JH, Turek M, Kang BG, Kwak JM, Kim WT (2012) Roles of four Arabidopsis U-Box E3 ubiquitin ligases in negative regulation of abscisic acid-mediated drought stress responses. Plant Physiol 160:556–568. 10.1104/pp.112.202143

Seo JS, Zhao P, Jung C, Chua N-H (2019) PLANT U-BOX PROTEIN 10 negatively regulates abscisic acid response in Arabidopsis. Appl Biol Chem 62:39. 10.1186/s13765-019-0446-0

Tang XL, Mu XM, Shao HB, Wang HY, Brestic M (2015) Global plant-responding mechanisms to salt stress: physiological and molecular levels and implications in biotechnology. Crit Rev Biotechnol 35:425–437. 10.3109/07388551.2014.889080

Trujillo M (2018) News from the PUB: plant U-box type E3 ubiquitin ligases. J Exp Bot 69:371–384. 10.1093/jxb/erx411

Trujillo M (2021) Ubiquitin signalling: controlling the message of surface immune receptors. New Phytol 231:47–53. 10.1111/nph.17360

Vierstra RD (2009) The ubiquitin-26S proteasome system at the nexus of plant biology. Nat Rev Mol Cell Biol 10:385–397. 10.1038/nrm2688

Wang N, Liu YP, Cong YH, Wang TT, Zhong XJ, Yang SP, Li Y, Gai JY (2016) Genome-wide identification of soybean U-Box E3 ubiquitin ligases and roles of GmPUB8 in negative regulation of drought stress response in Arabidopsis. Plant Cell Physiol 57:1189–1209. 10.1093/pcp/pcw068

Wang X, Ding YL, Li ZY, Shi YT, Wang JL, Hua J, Gong ZZ, Zhou JM, Yang SH (2019) PUB25 and PUB26 promote plant freezing tolerance by degrading the cold signaling negative regulator MYB15. Devel Cell 51:222–235. 10.1016/j.devcel.2019.08.008

Wiborg J, O’shea C, Skriver K (2008) Biochemical function of typical and variant Arabidopsis thaliana U-box E3 ubiquitin-protein ligases. Biochem J 413:447–457. 10.1042/BJ20071568

Wu Y, Wang W, Li Q, Zhang G, Zhao X, Li G, Li Y, Wang Y, Wang W (2020) The wheat E3 ligase TaPUB26 is a negative regulator in response to salt stress in transgenic Brachypodium distachyon. Plant Sci 294:110441. 10.1016/j.plantsci.2020.110441

Yan JQ, Wang J, Li QT, Hwang JR, Patterson C, Zhang H (2003) AtCHIP, a U-box-containing E3 ubiquitin ligase, plays a critical role in temperature stress tolerance in Arabidopsis. Plant Physiol 132:861–869. 10.1104/pp.103.020800

Yu F, Wu Y, Xie Q (2016) Ubiquitin–proteasome system in ABA signaling: from perception to action. Mol Plant 9:21–33. 10.1016/j.molp.2015.09.015

Zhang M, Zhao J, Li L, Gao Y, Zhao L, Patil SB, Fang J, Zhang W, Yang Y, Li M, Li X (2017) The Arabidopsis U-box E3 ubiquitin ligase PUB30 negatively regulates salt tolerance by facilitating BRI1 kinase inhibitor 1 (BKI1) degradation. Plant Cell Environ 40:2831–2843. 10.1111/pce.13064

Zhang YY, Yang CW, Li Y, Zheng NY, Chen H, Zhao QZ, Gao T, Guo HS, Xie Q (2007) SDIR1 is a RING finger E3 ligase that positively regulates stress-responsive abscisic acid signaling in Arabidopsis. Plant Cell 19:1912–1929. 10.1105/tpc.106.048488

Zhao JF, Zhao LL, Zhang M, Zafar SA, Fang JJ, Li M, Zhang WH, Li XY (2017) Arabidopsis E3 Ubiquitin Ligases PUB22 and PUB23 Negatively Regulate Drought Tolerance by Targeting ABA Receptor PYL9 for Degradation. Int J Mol Sci 18:1841. 10.3390/ijms18091841

